# Single-cell differential splicing analysis reveals high heterogeneity of liver tumor-infiltrating T cells

**DOI:** 10.1101/2020.03.22.002766

**Authors:** Shang Liu, Biaofeng Zhou, Liang Wu, Yan Sun, Jie Chen, Shiping Liu

**Author notes:** These authors contributed equally to this work as first authors (Shang Liu, Biaofeng Zhou). **Corresponding author:** Shiping Liu, BGI-Shenzhen, Beishan Industrial Zone, Shenzhen 518083, China.

## Abstract

Recent advances in single-cell RNA sequencing (scRNA-seq), enriched the knowledge of the heterogeneity of the tumor-infiltrating lymphocytes (TIL) for understanding the mechanisms of cancer initiation and progression. However, alternative splicing (AS), as one of the important regulatory factors of heterogeneity, has been poorly investigated. Here, we proposed a computational tool, DESJ-detection, which could fast and accurately detect the differentially expressed splicing junction (DESJ) between cell groups at single-cell level. We analyzed 5,063 T cells of hepatocellular carcinoma (HCC) and identified 1,176 DESJs across 11 T cell subtypes. Cell subtypes with a similar function clustered closer rather than the lineage at the AS level. Meanwhile, we identified two novel cell states, pre-exhaustion and pre-activation with the marker isoform CD103-201 and ARHGAP15-205. In summary, we presented a comprehensive investigation of alternative splicing differences, which provided novel insights for heterogeneity of T cells and can be applied in other full-length scRNA-seq datasets.

## 1. Introduction

T cell heterogeneity in the tumor microenvironment (TME) is tightly linked to tumor progress, prognosis, and therapies. The systematic interrogation of tumor-infiltrating lymphocytes has been fulfilled in liver[1], lung[2], colon[3] and breast cancers[4] using scRNA-seq. Effector and cytotoxic T cells can exert an anti-tumor effect by targeting tumor cells, and levels of effector CD8^+^ T cells are predictive of good survival in several cancers[5–7]. However, the tumor-infiltrating Tregs suppress the activity of T cell, myeloid cell, and stromal cells[8] by secreting immunosuppressive cytokines, such as FOXP3. Immunosuppressive cytokines then activate co-inhibitory receptors on T cells, such as PD1 and CTLA4, thus drives T cell dysfunction and exhaustion[9]. Meanwhile, the performance of these immunosuppressive cytokines and co -inhibitory receptors is influenced by alternative splicing. For example, one of the isoforms of *FOXP3* lacking exon 2 and exon 7 cannot perform the immunosuppressive function[10] and soluble *CTLA4* isoform shows the different effects on T cell state with full-length *CTLA4* isoform[11]. Therefore, Investigating the influence of alternative splicing on T cell state in TME will promote the understanding of T cell heterogeneity and the development of cancer therapy.

Alternative splicing analysis based on scRNA-seq is revolutionizing our understanding about the effect of alternative splicing on immune cells. Recently, scRNA-seq revealed the bimodality in splicing in immune cells while bulk RNA-seq might cover up the splicing difference between single cells[12]. However, the current computation framework in RNA-seq splicing analysis could not effectively detect the differential splicing between groups at the single-cell level. DEXSeq[13], rMATS[14], and MISO[15] were developed for bulk RNA-seq data. So, they might lead to incorrect results for their improper algorithms in single-cell transcriptome due to the low sequencing depth and high dropout rate. There were two programs were specially developed for scRNA-seq data, BRIE[16] and Outrigger[17]. But BRIE requires doing a pairwise comparison between every two cells to detect differential junctions, which is time-consuming and impractical. Outrigger utilizes the distribution mode of percent-spliced-in (Psi/J) to detect the differential splicing between cell groups. However, the distribution modes were just limited within five types and could not reflect the reality accurately. Thus, there is an urgent requirement to develop a convenient and effective computation tool to detect the differential splicing between groups.

To explore the T cell splicing heterogeneity in high resolution, we have developed a novel computation framework, DESJ-detection, to detect differential splicing between groups at the single-cell level. We applied it to a published scRNA-seq dataset from HCC patients. We identified 1,176 DESJs across the 11 cell clusters and found the functional similar T cell subsets shared a similar splicing pattern. We revealed the relationship between alternative splicing and T cell functional subpopulations, especially pre-exhausted and pre-activation subpopulations. Thus, the systematic evaluation of differential splicing across T cells in TME of HCC provides comprehensive knowledge of the alternative splicing characteristics of TILs and will facilitate the progress of cancer diagnosis and treatment.

## 2. Material and methods

### 2.1 Data Sets

We downloaded the scRNA-seq raw reads of Human T cells in Fastq from EGD database (EGAS00001002072). The corresponding gene expression matrix was downloaded from GEO database (GSE98638). This dataset contained 5,063 T cells assigned into 12 clusters[1]. These T cells were sampled from peripheral blood, tumor, and adjacent normal liver tissue. The detailed clinical information of patients and cell clusters information was listed in Table 1. The human genome with the version of GRCH38 was taken as the alignment reference using by STAR[18].

**Table 1.** Annotation about cell clusters

| Cluster | Cell number | Function annotation | Type |
| --- | --- | --- | --- |
| C01_CD8.LEF1 | 161 | Naïve T cell | CD8+ T cell |
| C02_CD8.CX3CR1 | 288 | Effector T cell | CD8+ T cell |
| C03_CD8.SLC4A10 | 363 | MAIT | CD8+ T cell |
| C04_CD8.LAYN | 300 | Exhausted T cell | CD8+ T cell |
| C05_CD8.GZMK | 467 | T cell in mediate state | CD8+ T cell |
| C06_CD4.CCR7 | 646 | Naïve T cell | CD4+ T cell |
| C07_CD4.FOXP3 | 261 | Peripheral Treg | CD4+ T cell |
| C08_CD4.CTLA4 | 582 | Tumor Treg | CD4+ T cell |
| C09_CD4.GZMA | 689 | T cell in mediate state | CD4+ T cell |
| C10_CD4.CXCL13 | 146 | Exhausted T cell | CD4+ T cell |
| C11_CD4.GNLY | 167 | Effector T cell | CD4+ T cell |
| Unknown | 993 | NA | NA |

### 2.1 Pipeline for creation of junction count matrix

we used the developed pipeline to create the junction count matrix. Firstly, we merged all the SJ.out.tab files output by the STAR aligner. Secondly, we retained junctions that were detected more than R_m_ reads in at least Cell_m_ cells (Cell_m_ = 10, R_m_ = 4, by default). Thirdly, we only retained the junctions that unique annotated by one gene. At last, we obtained the count matrix containing the read number of junctions in each cell (Figure S1 A).

### 2.2. Description of software to detect differential junction usage

The software required four inputs: junction count matrix (matrix A), junction annotation file (from the pipeline we developed), the uniquely mapped read number of each cell, and cell clustering information. For a schematic illustration of differential splicing analysis process, refer to Figure 1 A. Firstly, we extracted junctions of a single gene (Gene1) from matrix A and normalized it with the number of uniquely mapped reads to obtain matrix C. Then, we performed iteration k-means for cells in matrix C to outlier the cells (SD < 0.2 and Mean < 1 by default) (precise steps are shown Algorithm 1). Third, we normalized the remained cells with all the junction reads count of Gene1 (matrix D). Finally, we used limma-trend to detect the DESJs between groups. The software output a res.xls file including statistically significant DESJs (adj.p.value ≤ 0.01 and log2(FC)≥ 1 or ≤ −1) and junction expression heatmaps of each gene with DESJs.

**Figure 1.**
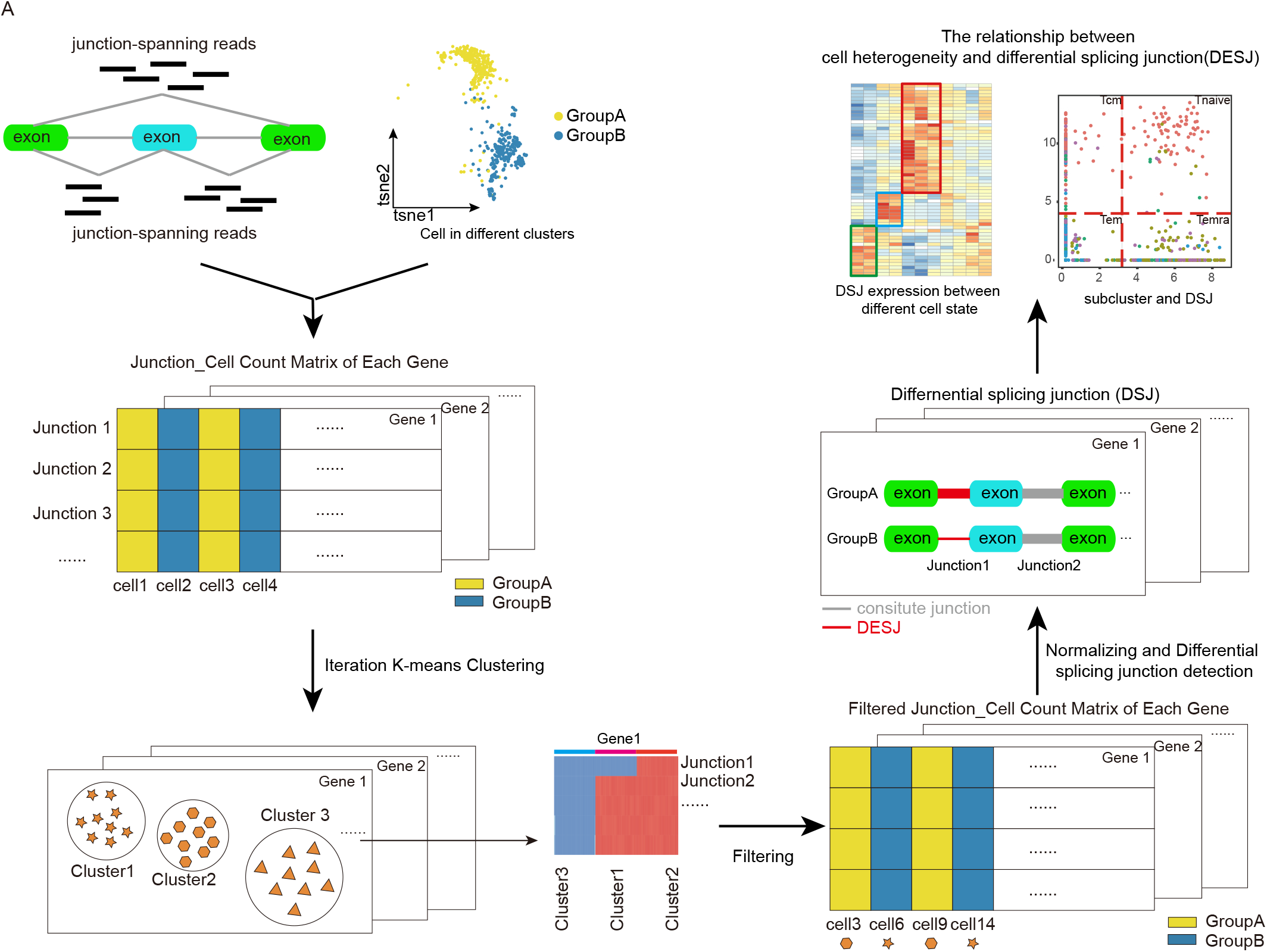
The overview of DESJ-detection.

#### Algorithm 1: Filter outlier

~~~
Input: Matrix C, maxsd, maxmean
Output: Outlier cells list
two_clusters ← kmeans(Matrix C)
cell_list ← Find the cell list with minimum mean and standard deviation comparing to the other cluster
Matrixtmp ← Matrix C[,cell_list]
Meantmp ← rowMeans(Matrixtmp)
Sdtmp ← rowSd(Matrixtmp)
while Meantmp < maxmean & Sdtmp < maxsd do
      two_cluster ← kmeans(Matrixtmp)
      cell_list ← Find the cells with minimum mean and sd comparing to the other clusters
      Matrixtmp ← Matrix C[,cell_list]
      Meantmp ← rowMeans(Matrixtmp)
      Sdtmp ← rowSd(Matrixtmp)
return cell_list
~~~

### 2.3 Simulate scRNA-seq data

We adopted a simulating strategy to evaluate the performance of our method. Firstly, we selected 200 genes from human GTF files among which 100 genes would be simulated as containing differential alternative splicing events. Then we further selected two transcripts for each gene to get a total of 400 transcripts. Next, we simulated RPK (reads per kilobase) value of 400 transcripts by a Perl script. RPK ratio of two transcripts belongs to the same gene was reciprocal between two cells while the RPK value of gene was a constant, such as 25, 50, 100, and 200. Two simulated cells were labeled as group A and group B, respectively. The cells with the same label were simulated a similar RPK ratio of two transcripts belong to the same gene. In addition, we set four levels of log2(RPK ratio) as 0,1,2,3 to simulate the degree of junction differential expressed. Besides, we stimulated the dropout ratio as four levels: 0, 0.1, 0.2, 0.4 by applying the simulator strategy of BRIE. Finally, fastq files were generated by Spanki[19] with errorfree mode and splicing junction information was acquired. We obtained 200 cells for each condition. The description of simulation was displayed as follows. Visit GitHub for more detailed information. (https://github.com/lucky-Mendel/DSJ-detection-simulator)

1. Obtain sim_rpk files for each pair of cells using a Perl script.
2. (necessary if dropout >0) generate a dice format file as input in step3.
3. (necessary if dropout >0) simulate dropout event with a modified script coming from the BRIE simulator.
4. Output fastq files by Spanki.

### 2.4 Differential gene expression analysis and gene set enrichment analysis

We performed the Limmar R package to analyze differential expressed genes between two target clusters. The significant genes were identified as those met these criteria: 1) FDR adjusted p-value of F test < 0.01; 2) the absolute value of log2 fold change was larger than 2. After differential gene expression analysis, we obtained the genes which were highly expressed in one group. We performed gene set enrichment analysis[20, 21] by the web-based tool provided by broad institute (http://software.broadinstitute.org/).

### 2.5 Survival analysis

The TCGA LIHC data were applied to assess the relationship between patient survival and individual genes, individual isoforms, and gene sets from specific cell clusters. We downloaded the data of gene expression and isoform count from UCSC Xena[22] (http://xena.ucsc.edu/) and retrieved clinical data from the Genomic Data Commons Data Portal (https://gdc-portal.nci.nih.gov/). Three hundred and seventy-seven patients without immunotherapy treatment were included in the survival analysis. Firstly, the isoform read count data were normalized by the isoform’s length and uniquely mapped read number of each patient. Then. to rectify the influence of T cell compositions within each sample, the expression of selected genes and isoforms in the tumor were divided by the geometric mean expression of *CD3* genes. *CD3* gene expression was assigned as the arithmetic mean of the corresponding isoforms (*CD3D*, *CD3E* and *CD3G*). Thirdly, for each selected genes and isoforms, we set the relative expression lower and upper threshold as the median minus or plus 10% MAD (median absolute deviation) respectively. Fourthly, we retained the samples whose relative expression is beyond these thresholds then divided patients into high and low expression groups. To explore whether CD103-201 isoform was correlated with prognosis, we calculated two scores for each patient by a weighted sum of fold change value of signature genes between CD 103-201^+^ population and CD103-201^-^ (Supplementary Table 1) and gene expression in TCGA data. Then, the patients were split into two groups by the median value of the expression score of patients. The statistical analysis was performed using the R package “survival”.

### 2.6 Developmental trajectory inference

We used the Monocle (version 2)[23] to order CD8/CD4 T cells in pseudo time respectively. TPM value was converted into normalized mRNA counts by the “relative2abs” function in monocle then we created an object with the parameter “expressionFamily = negbinomial.size”. Finally, the CD8^+^/CD4^+^ T cell differentiation trajectory was determined by the default parameters of Monocle.

### 2.7 Definition of exhaustion and naïveness scores

Similar to Guo et.al[2], we firstly identified the most significant genes between exhausted T cluster (C04_CD8.LAYN) and other T clusters using moderated t-test implemented by the R package limma (log2(FC) >= 4 & FDR < 0.01). Then, we defined the exhaustion score for CD8^+^ T cells as the average expression of these markers after z-score transformation (original value is log2(TPM+1)). A similar method was used to define naïveness scores for CD4^+^ T cells using the common naïve markers. Finally, we calculated the significant level of the exhaustion and naiveness scores of cells from different clusters by t-test.

## 3 Results

### 3.1 The overview of DESJ-detection

Revealing splicing differences at the single-cell level would deepen our understanding of cell heterogeneity, function, and phenotype. The major challenges in differential splicing analysis are scRNA-seq data has much dropout events and low sequencing depth compared to bulk RNA-Seq, which hinders reflection of the real splicing structure of genes. Besides, splicing analysis in RNA-Seq data mainly is limited in SE (exon skip events) and MXE (mutually exclusive exon events). To address these two challenges, we proposed DESJ-detection, an algorithm that uses junction-spanning reads to detect DESJs. Firstly, we input all the junctions read count of each cell and output junction-cell count matrix of each gene. Secondly, we applied iterative K-means to cluster cells and removed the clusters with low expression (SD < 0.2 and Mean < 1) of all junctions resulting from low coverage and high dropout rate. Then, we utilized a new normalization method to eliminate the influence of differential expression at the gene level on differential junction expression detection. Specifically, it normalized the junction read count with the read count of each gene rather than uniquely mapped reads of each cell. Finally, we identified DESJs based Limma-tread algorithm with the value of fold change (FD) and adjusted p-value. Meanwhile, DESJ-detection can detect the DESJs at any region of a gene, so it can discover any patterns of alternative splicing, rather limited in SE and MXE events (Fig. 1A). We also developed a convenient pipeline (https://github.com/liushang17/DESJ-detection), which starts from the generation of junctions, filtering and annotation of junctions, preparation of junction count table and detection of DESJs (Fig. S1A).

To assess the performance of the software in differential alternative splicing detection, we simulated scRNA-seq data with a pipeline based Spanki considering different factors, including reads coverage, dropout rate, and degree of junction differential expressed. Our method was proved to be effective. For example, the simulated cells were divided into five clusters by the expression of two isoforms of PPT1. As we can see, four cell clusters showed junction differential expression and another cluster with low gene expression was removed by the iterative K-means clustering, because it failed to reveal the real junction usage (Fig. S1B). In the meantime, we observed the sensitivity level reached up to about 70% even at the lowest coverage level (RPK = 25) when the junction differential expression is more than control and without dropout events. And the sensitivity was essentially maintained at 85% at the general coverage level (RPK>=50). In addition, the sensitivity also reached a high percentage (>=70%) when dropout rate is more than 0 (Fig. S1 C). Besides, more than 95% of identified genes was DESJ related genes. Taken together, DESJ-detection proved its robustness in dropout events, low coverage requirement for detection, and high sensitivity to DESJs.

### 3.2 Differential usage of junctions in UTR regions across T cell clusters

we performed DESJ-detection in a published scRNA-seq data set. It includes 5,063 T cells from tumor tissues, normal tissues, and peripheral blood of six HCC patients and had been assigned into 11 T cell subsets, including naïve T cells (C01_CD8.LEF1, C06_CD4.CCR7), effector T cells (C02_CD8.CX3CR1, C11_CD4.GNLY), exhausted T cells (C04_CD8.LAYN, C10_CD4.CXCL13), Tregs (C07_CD4.FOXP3, C08_CD4.CTLA4), mucosal-associated invariant T cells (C03_CD8.SLC4A10), and intermediate T cells (C05_CD8.GZMK, C09_CD4.GZMA). We obtained a set of 134,414 junctions that satisfied read count more than 4 in at least 10 cells, covering 12,587 genes (Fig. S2 A & Fig. S2 B). We further filtered the junctions that not located at any annotated genes or located at fusion genes. In the end, we retained 119,311 junctions from 10,556 genes. By DESJs analysis, we finally identified 1,176 DESJs across 11 clusters (log2(FC) ≥ 1, adjust p-value ≤ 0.01) (Supplementary Table 2).

To characterize the distribution of DESJs in genomics, we investigated the frequency of DESJs between different genome regions. We found significant higher frequency of DESJs in UTR regions than coding regions between clusters (p-value = 0.004 for CD8^+^ T cells and p-value = 6.456e-13 for CD4^+^ T cells, Student’s t-test. Fig. 2 A & Fig. S2 C & Fig. S2 D). This may result from longer junction length (end site – start site + 1) in UTR region. We additionally observed that DESJs are significantly longer than nonDESJ both in UTR and Coding regions. Besides, the DESJs in UTR regions were also longer than the these in Coding regions (Fig. 2B). Because UTR regions are longer than coding regions, these two phenomena may be explained that the longer junctions would provide more possible splice sites and potential regulation functions. Previous studies have revealed that the ratio of genes whose UTR region happened alternative splicing made up to 10%-18% [24, 25]. Besides, alternative splicing in UTR also made a great contribution to regulate gene expression[26]. Therefore, our results revealed that alternative splicing in UTR region has been underestimated because of its higher frequency of differential alternative splicing between cell clusters. Specifically, we supposed alternative splicing in UTR regions may contribute a lot to not only gene expression regulation but also cell heterogeneity.

**Figure 2.**
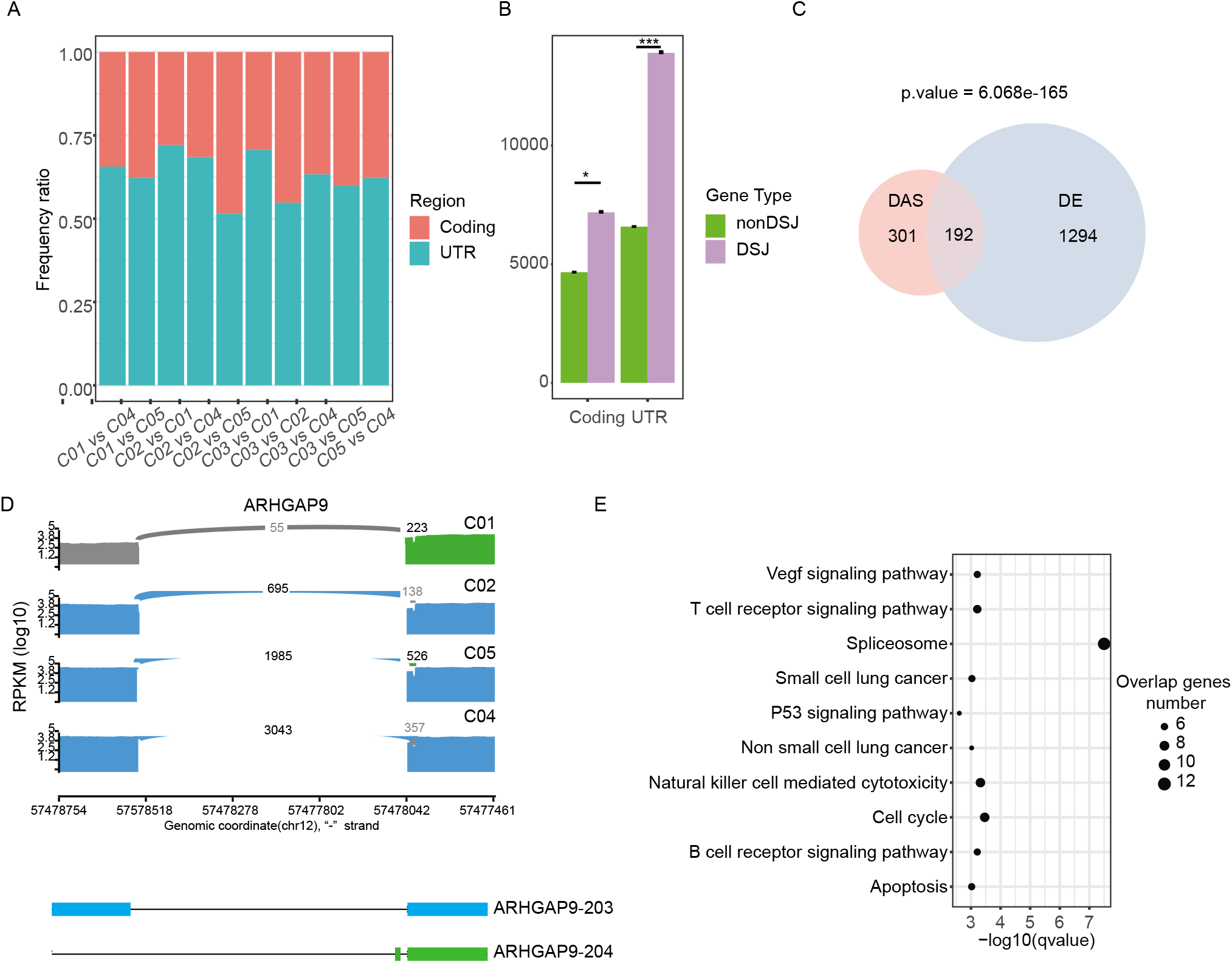
Wide differential usage of junctions in the UTR regions across T cell clusters. **A,** The proportions of differential splicing junction frequency in the UTR regions and the Coding regions across CD8+ T cells. **B**, The length difference between DESJ and nonDESJ in Coding region and UTR region. **C**, The overlap between differential expression genes and DESJ associated genes among T cells clusters. **D**, Sashimi plots illustrating the read distribution of ARHGAP9 in CD8+ T cells from P0508. The colors represent different isoforms. This alternative splicing of ARHGAP9 happens in the UTR regions. Naïve T cells (C01_CD8.LEF1) show obvious differential usage of isoforms from other clusters. **E**, Result of KEGG pathway analysis for genes with differential splicing in the UTR regions.

At the meantime, we noticed that the considerable differential expression genes between clusters were also DESJ related genes in T cells (Fig. 2 C). In addition, there were about 60% differential expression genes possessing DESJs in the UTR region (Fig. S2 G). For example, *ARHGAP9,* a member of RhoGAP family and associated with good survival, was a differential expressed gene (highly expressed in C04_CD8.LAY, C10_CD4.CXCL13, and C08_CD4. CTLA4). And it also showed differential splicing in UTR region between CD8^+^ T cell clusters (C01_CD8.LEF1, C02_CD8.CX3CR1, C04_CD8.LAYN, C05_CD8.GZMK) (Fig. 2 D & Fig. S2 F). Specifically, ARHGAP9-203 was upregulated in exhausted T cells and Tumor-infiltrated Tregs, while ARHGAP9-204 was mainly expressed in naïve T cells and peripheral blood Tregs (Fig. 2 D).

KEGG pathway analysis of genes possessing DESJs in UTR was mainly involved in the VEGF signaling pathway, T cell receptor signaling pathway, spliceosome, P53 signaling pathway, and cell apoptosis. Meanwhile, the genes with DESJs in the Coding region were associated with innate immune pathways and spliceosomes. Hence, these emphasized the alternative splicing in UTR regions may relate with the specific function of cells. Taken together, our results indicated that alternative splicing in UTR regions may play a regulated role in gene expression between cell clusters.

### 3.3 T cell heterogeneity at splicing level

To explore the association between alternative splicing and the heterogeneity of T cell function, we further utilized the identified DESJs across T cell clusters to obtain cell-type-specific splicing junctions. In this study, we detected 335 DESJs from 165 genes among CD8^+^ sub-clusters and 484 junctions from 239 genes among CD4^+^ sub-clusters (Supplementary Table 2). We used two distinct datasets to hierarchically clustered T cells, the number of DESJs and the expression of DESJs across all cell clusters. Both of them indicated that cells with a similar function rather than the lineage, exhibited a similar alternative splicing pattern. (Fig. 3 A). For example, tumor-infiltrating Treg and exhausted T cell (C04_CD8.LAYN, C08_CD4. CTLA4, C10_CD4.CXCL13) clustered together, demonstrating a huge difference between them and others. What’s more, naïve T cell (C01_CD8.LEF1, C06_CD4.CCR7), effector T cell (C02_CD8.CX3CR1, C11_CD4.GNLY), and intermediate state (C05_CD8.GZMK, C09_CD4.GZMA) clustered together respectively. In the meantime, we mentioned that exhausted T cells showed the most significant difference in the DESJ number with other T cells, indicating exhausted T cells would emerge the greatest changes in alternative splicing. These results indicated junction usage difference between cell clusters mainly depends on the functional state of the clusters.

**Figure 3.**
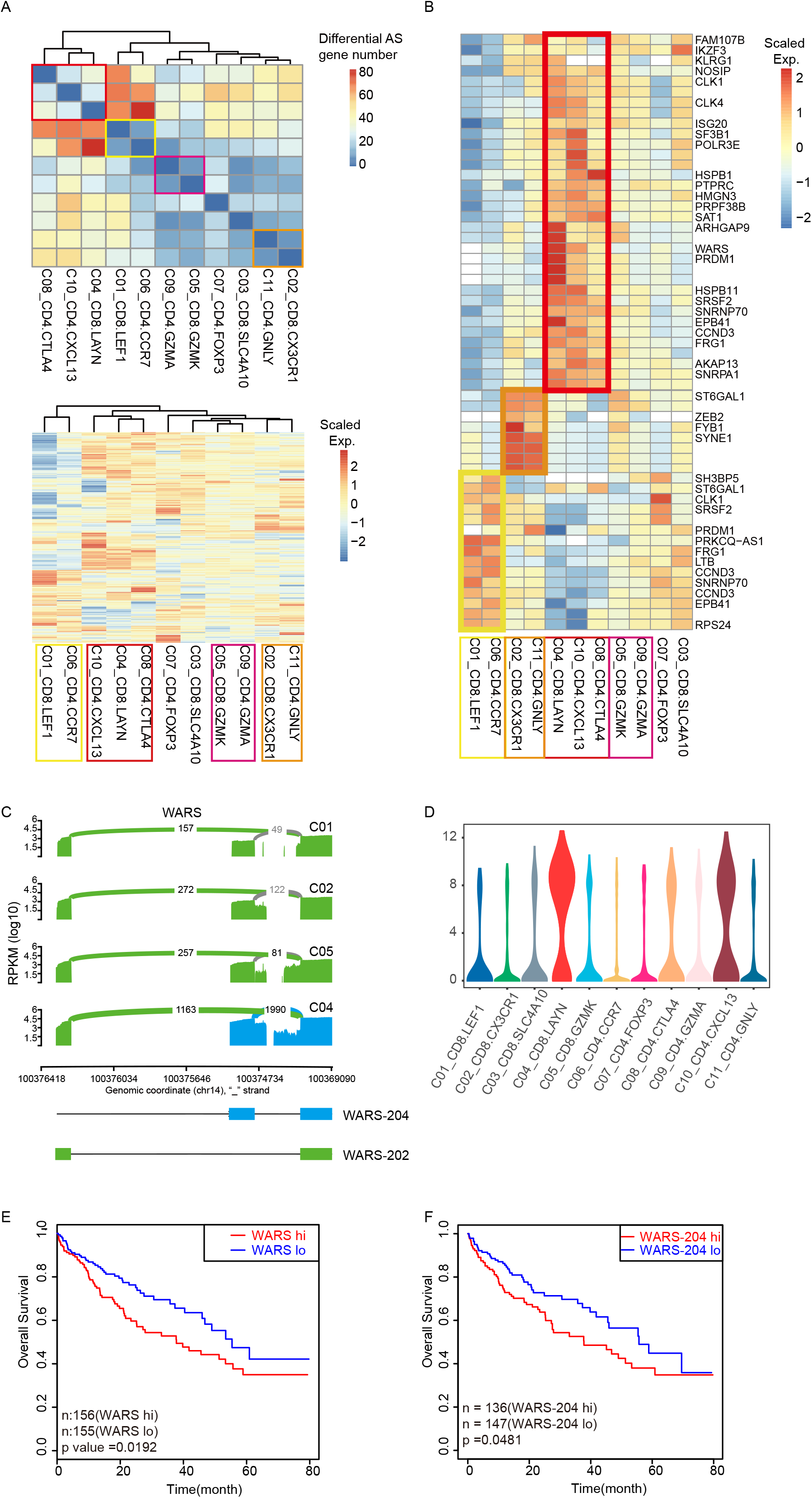
T cell heterogeneity at splicing level. **A,** Upper: Heatmap of DESJ associated genes number between pairwise clusters across T cells. Below: z-score normalized mean expression of all DESJ in each T cell cluster. Boxes with different colors highlight the patterns of different functional T subtypes. **B**, the z-score normalized mean expression of selected DESJ that shares in the similar function subtypes. **C**, Violin plots comparing the expression of WARS in 11 T cell clusters. **D**, The disease-free survival curve based on TCGA HCC data showing patients with higher expression of WARS had a poor prognosis. **E**, Sashimi plots illustrating the read distribution of WARS in CD8+ T cells from P0508 patient. WARS-204 highly expresses in exhausted T clusters (C04_CD8.LAYN). **F**, Disease-free-survival (DFS) curve comparing the high and low expression of WARS-204 based on the TCGA HCC cohort shows higher expression of WARS-204 in tumor means bad prognosis.

We next focused on the DESJs related genes between four functional states, including naïve T cells, effector T cells, exhausted T cells, and mediate T cells. Naïve and exhausted T cells mainly showed differential splicing in genes of splicing and immune, such as *CD45, HSPB1, CLK1, SRSF2, SNRNP70.* Effector T cells were characterized by the differential splicing in *ZEB2, FYB1,* and *SYNE1* (Fig. 3 B). Among all the DESJs related gene, *WARS* that highly expressed in exhausted T cell and being a maker of exhaustion, showed differential splicing between exhausted T cell and other T cells (Fig 3 C, D & Fig. S3 A). The junction (chr14_100369259_100376259_2), representing WARS-202 showed the widespread expression in all T cells while the junction (chr14_100369259_100375282_2), representing WARS-204, only widely expressed in Tregs (C08_CD4. CTLA4) and exhausted T cells (C04_CD8.LAYN, C10_CD4.CXCL13). Prognostic analysis with TCGA LIHC data revealed that the upregulated expression of WARS was associated with worse prognosis (Fig. 3 E). Thus, we proposed the upregulation expression of isoform WARS-204 actually represented worse prognosis (Fig. 3 F). Furthermore, the prognostic analysis with TCGA LIHC data at the isoform level confirmed our hypothesis (Fig. S3 B).

This case inspirited us whether some immunity therapy-related target genes also displayed such a phenomenon. We found two T cell immunity checkpoint genes whose upregulation expression related to worse prognosis, *TNFRSF4,* and *HAVCR2*. However, only one of its isoforms is in accordance with its gene performance, implying that this isoform may be as an actual therapy target (Fig. S3 C). In summary, these results demonstrated alternative splicing would have a huge effect on the function and phenotype of T cells and would be potential markers for cancer prognosis and treatment.

### 3.4 Two novel functional subpopulations identified by CD103-201 and ARHGAP15-205

To further show the inner heterogeneity in T cell clusters, we utilized DESJs to identify the functional subpopulations. After that, we inferred the potential function of isoforms. *ITGAE,* also known as *CD103,* is a tissue-resident T cell marker and highly expressed in exhausted T cells. One of its isoforms, CD103-201, was upregulated in CD8^+^ exhausted T cells (CD4_CD8.LAYN), while another isoform CD103-202 universally expressed in all CD8^+^ T cells. The differential splicing of *CD103* was not any common alternative splicing patterns (SE, MXE, IR, A5SS, and A3SS), resulting from CD103-201 expressed nine more exons than CD103-202 (Fig. 4 A). In addition, we observed CD103-202 showed widespread expression in C04_CD8.LAYN, while CD103-201 exhibited a bimodal pattern in C04_CD8.LAYN, implying that different isoforms of *ITGAE* would play different roles in exhausted T cells function (Fig. 4 B). Meanwhile, the two populations (CD103-201^+^ and CD103-201^-^ populations) displayed obviously different expression patterns. Specifically, CD103-201^+^ population highly expressed exhausted marker *ENTPD1,* while CD103-201^-^ population showed high expression of ribosome proteins, including *RPL27, RPL35A, RPS29, RPS21* (Fig. 4 C). GO enrichment analysis shows CD103-201^-^ population own extremely strong translation vitality, indicating CD103-201^-^ population may be in the state of transition (Fig. S4 A). To verify this hypothesis, we performed pseudotime analysis among all the four CD8^+^ clusters. CD103-201^-^ population and C05_CD8.GZMK were located more centrally in the Monocle trajectory and had a significantly lower exhaustion score than CD103-201^+^ population. Thus, this result further suggested that CD103-201^-^ population was possible in “pre-exhaustion” state (Fig. 4 D). As expected, we also observed that CD103-201^-^ population was associated with better prognosis in TCGA LIHC data compared with CD103-201^+^ population (P= 0.009, Cox regression, Fig. S4 B). In summary, these results represented CD103-201 may be associated with T cell exhaustion and have the potential to clinical application.

**Figure 4.**
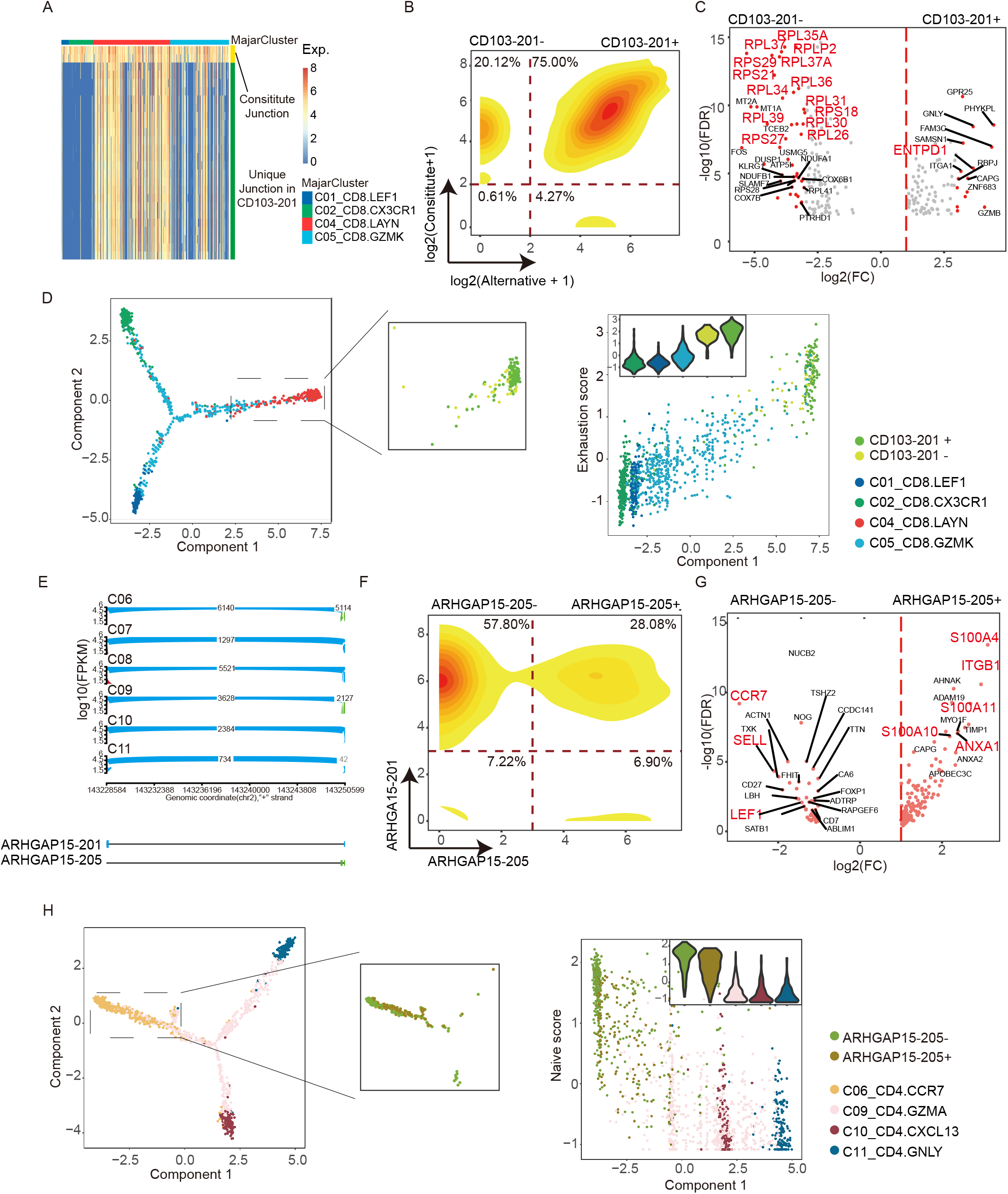
Two novel functional sub-population identified by CD103-201 and ARHGAP15-205. **A,** Heatmap of 4 CD8^+^ clusters with junctions from *ITGAE (CD 103).* Alternative junctions, representing CD103-201, highly express in exhausted T cells (CO4_CD8.LAYN), while constitute junctions universally express in all T cell clusters. **B**, The bimodal distribution of alternative junctions from *CD103* shows the Intrinsic heterogeneity in exhausted T cells (CO4_CD8.LAYN). Cell density is color-coded, with red denoting high density and yellow low density. **C**, Volcano plot showing differentially expressed genes between the CD103-201^+^ and CD103-201-populations. Each red dot denotes an individual gene with adjusted P-value < 0.01 (two-sided moderated t-test with limma) and abs(fold change) ≥ 8. **D**, Result of GO enrichment analysis for genes highly expressing in CD103-201- population. **E**, Disease-free-survival (DFS) curve comparing the high and low expression of marker genes of CD103-201^+^ high population and CD103-201- population. **F**, Sashimi plots illustrating the read distribution of *ARHGAP15* in CD4^+^ T cells from P0508. ARHGAP15-205 highly expresses in naïve T cells (CO6_CD4.CCR7). **G**, Similar to *CD103,* the bimodal distribution of ARHGAP15-205 shows the Intrinsic heterogeneity in naive T cells (CO6_CD4.CCR7). **H**, The Venn graph showing the overlap of activation associated genes identified in this study with those from previous studies by De Simone et al. (2016) (p = 2e-85) determined by hypergeometric test. **I**, Volcano plot showing differentially expressed genes between the ARHGAP15-205+ and ARHGAP15-205-populations. Each red dot denotes an individual gene with adjusted P value < 0.01 (two-sided moderated t-test with limma) and abs(fold change) ≥ 2. **J**, Result of GO enrichment analysis for genes highly expressing in ARHGAP15-205+ population.

*ARHGAP15,* a Rac1-specific GAP, was reported to be associated with the development of diverse tumors, including colorectal cancer[27], glioma[28] and pancreatic ductal adenocarcinoma[29]. However, it is little-known about the relationship between T cell state and *ARHGAP15* at the isoform level. Our study discovered that ARHGAP15-201 expressed universally in all cell clusters, but ARHGAP15-205 exhibited a highly specific expression pattern (Fig. 4E). Further, ARHGAP15-205 shows a striking bimodal expression distribution in both CD8 naïve T cells (C1_CD8-LEF1) and CD4 naïve T cell (C6_CD4-CCR7) (Fig. 4F & Fig. S4 E). This implied ARHGAP15-205 may affect the functional state of naïve T cell. We identified 174 genes highly expressed in ARHGAP15-205^+^ naïve T cell (FDR < 0.01, log2(Fold change) ≥ 1) (Supplementary Table 1). These genes significantly overlapped with signature genes of cell cluster in the activated state, which was identified by three previous studies (Fig. S4 C). Thus, we supposed the ARHGAP15-205^+^ population has a similar activation characteristic. Signature genes of ARHGAP15-205^+^ include *S100A4, ITGB1, S100A6,* and *LGALS1,* supporting that ARHGAP15-205^+^ population trend toward activation state (Fig. 4 G). On the contrary, ARHGAP15-205^-^ population highly expressed the genes related to resting state, including *CCR7, SELL* and *LEF1,* demonstrating it is in the relative resting state (Fig. 4 G). Meanwhile, GO biological process enrichment analysis showed ARHGAP15-205^+^ population signature genes enriched in the cell differentiation (including leukocyte differentiation and lymphocyte differentiation) and cell activation (Fig. S4 D). What’s more, the pseudotime analysis of cells in C06_CD4.CCR7, C09_CD4.GZMA, C10_CD4.CXCL13, and C11_CD4.GNLY showed ARHGAP15-205^+^ cells were closer to cells in C09_CD4.GZMA and had a lower naïve score compared with ARHGAP15-205^-^ population (Fig. 4 H). These results suggested that ARHGAP15-205^+^ CD4 naive T cells, might be in the pre-activation state and possess immune killing function. The identical performance also emerged in CD8 naïve T cell (C01_CD8-LEF1) (Fig. S4 E, F). Altogether, these results emphasized that alternative splicing analysis at single-cell level would reveal cell heterogeneity and discover cell sub-clusters in higher resolution than gene expression level.

## 4 Discussion

The scRNA-seq technology has developed rapidly and has been widely applied in many frontier fields including tumor heterogeneity, cell differentiation, and neural development. Compared to 3’ enrichment methods, full-length single-cell RNA data can not only quantify gene expression but also analyze the structure of genes in high resolution, such as single nucleotide variants (SNV) and alternative splicing (AS) detection. Due to the lack of available software to analyze cell heterogeneity with alternative splicing, single-cell research currently is still limited to gene expression profiling. Here we have developed a differential alternative splicing detection software for the full-length scRNA-seq dataset.

DESJ-detection was proved to detect the DESJs between different cell types at the single-cell level in a robust and effective way. To detect the DESJs between two cell groups would be affected by the technical limitation in scRNA-Seq, which would lead to the incorrect representation of the alternative splicing structure. The iteration k-means could effectively find and filter these cells. It works like wringing out two sponges containing water, only when as much as water was removed would the properties of the two sponges themselves be compared accurately. In addition, the different read count of genes, resulting from different sequence depth and gene expression level, would prevent the precision detection of DESJs. Taking these two factors into consideration, we normalized the junction matrix of each gene with reads count of the gene in each cell. This computation framework suggested a novel strategy to detect the differential alternative splicing between two groups of cells at the single-cell level and filled the gap in this field. However, DESJ-detection could not accurately detect the isoforms composition of a single cell for any given genes because some junctions may not uniquely belong to one isoform. Additional work to develop an improved version to address the above shortcoming is ongoing and would result in the interpretation of isoform difference in higher resolution.

We performed DESJs-detection in a T cell dataset from six patients diagnosed with HCC provides novel insight into T cell heterogeneity. One interesting finding is cell clusters with a similar functions displayed a minor number of DESJs related genes comparing with others and possessed a similar DESJs expression pattern. For example, exhausted T cells and tumor-infiltrating Tregs, which shared similar high expression genes *LAYN, HAVCR9,* and *ENTPD1,* also shared similar splicing patterns, such as *WARS, ARHGAP9, SRSF2.* These relationships may partly be explained that cells with a similar function would share similar expression profiles of genes as well as isoforms. At the same time, some unique isoforms in exhausted T cells are related to poor prognoses, such as *WARS* and *CCND3*. Therefore, altering the isoform preference of specific genes in T cells might be another way for cancer immunotherapy.

The association between alternative splicing and the cell clusters may be applied to infer the function of alternative splicing and predict novel subpopulations. For example, CD103-201 revealed a novel sub-cluster, pre-exhausted population. And then, CD103-201 may be inferred to play a role in T cell exhaustion in liver cancer. A similar phenomenon emerged at ARHGAP15-205, an isoform related to T cell activation. Further studies are needed to affirm these results by experiments and interrogate the potential mechanism, as well as other isoforms related to the cell functional state.

With the rapid development of scRNA-seq, smart-seq3[30] with longer read length and faster sequencing has emerged, leading to researches on single-cell alternative splicing a hot topic. However, the conditions to support single cell alternative splicing analysis, including sequencing depth and coverage have not been explicitly disclosed. Secondly, there is still a lack of corresponding methods on how to construct a profile of alternative splicing at the single-cell level. Finally, the combined analysis of single-cell alternative splicing and gene expression has not been explored. Our program will greatly contribute to enriching the research strategy of alternative splicing and exploring its potential function extensively.

## Authors’ contributions

All authors contributed to the study conception and design. Material preparation, data collection was done by Yan Sun and Jie Chen. Analysis were performed by Shang Liu and Biaofeng Zhou. The first draft of the manuscript was written by Biaofeng Zhou and Shang Liu. All authors commented on previous versions of the manuscript. All authors read and approved the final manuscript.

## Consent for publication

All the authors agreed to publish the work.

## Ethics approval and consent to participate

Not applicable.

## Acknowledgments

This study is supported by Science, Technology and Innovation Commission of Shenzhen Municipality under grants (No. GJHZ20180419190827179 and No. JCYJ2017041253248372). We also thank Peking University BIOPIC Data Access Committee (PUBDAC) for approving downloading raw single-cell RNA data of HCC as well as Si Qi for her proper guide about article writing.

## Availability of data and materials

RNA-seq data of human T cells in fastq format was downloaded from EGD database with accession study title EGAS00001002072. Matrix of gene expression was downloaded from GEO database with accession numbers GSE98638. Analysis code of such HCC data can also be found at https://github.com/liushang17/DESJ-detection.

---

**Figure S1.**
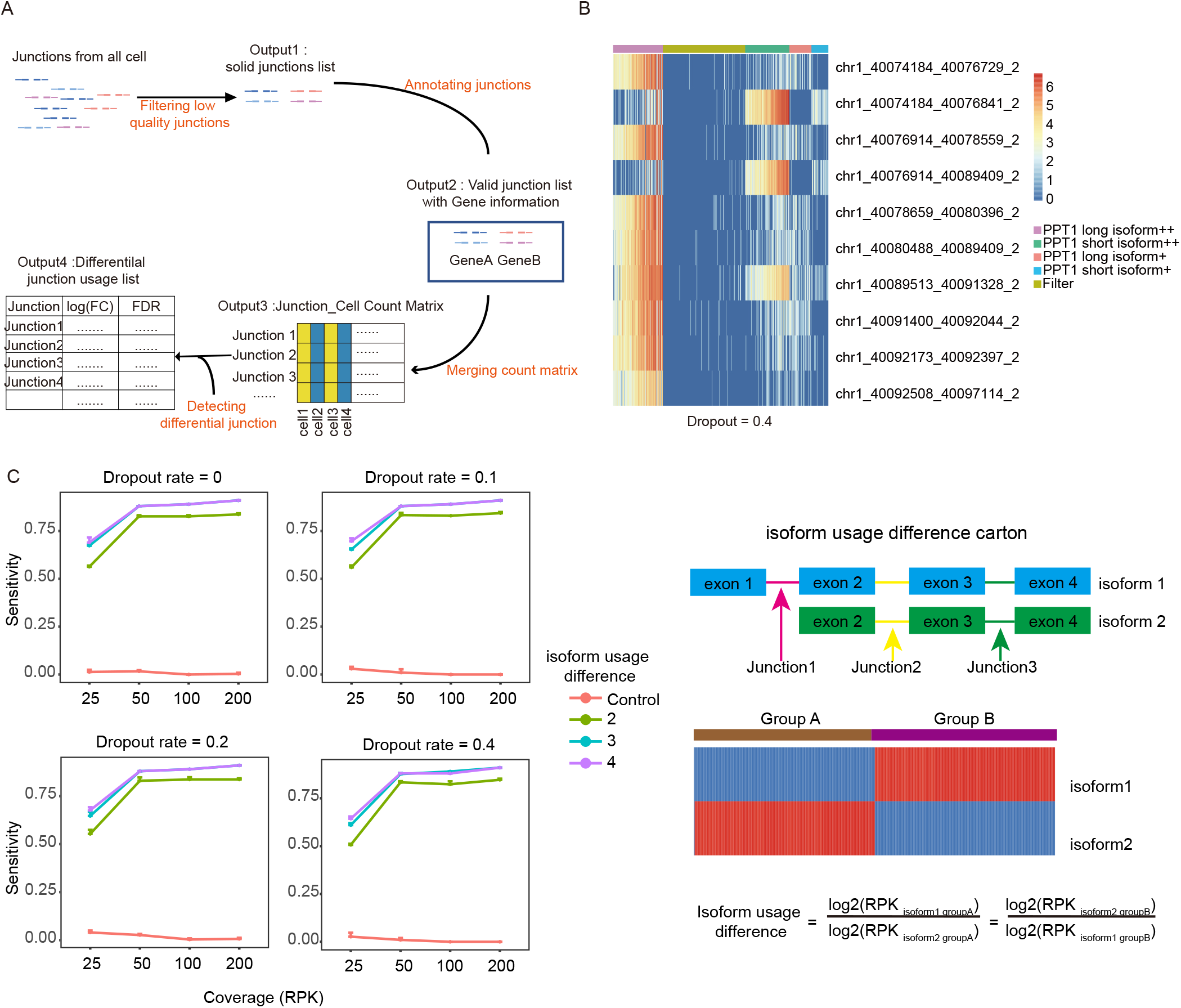
The diagrammatic sketch of the pipeline for the creation of junction count matrix and the performance of DESJ-detection on simulated data. **A,** The preparation work (see method for detailed information). **B**, Heatmap of five identified clusters by iterative K-means with all junction of PPT1 from simulated data. The Violet bar represents cells highly expressing PPT1 long isoform. The green bar represents cells highly expressing PPT1 short isoform. The pink bar represents a cell moderately expressing PPT1 long isoform. Blue bar represents cell moderately expressing PPT1 short isoform. Yellowish-brown bar represent cell failing to reflect the composition of PPT1 isoforms. **C**, Left shows the sensitivity of DESJ detection by DESJ-detection. Right is the schematic diagram of isoform usage difference of simulated data.

**Figure S2.**
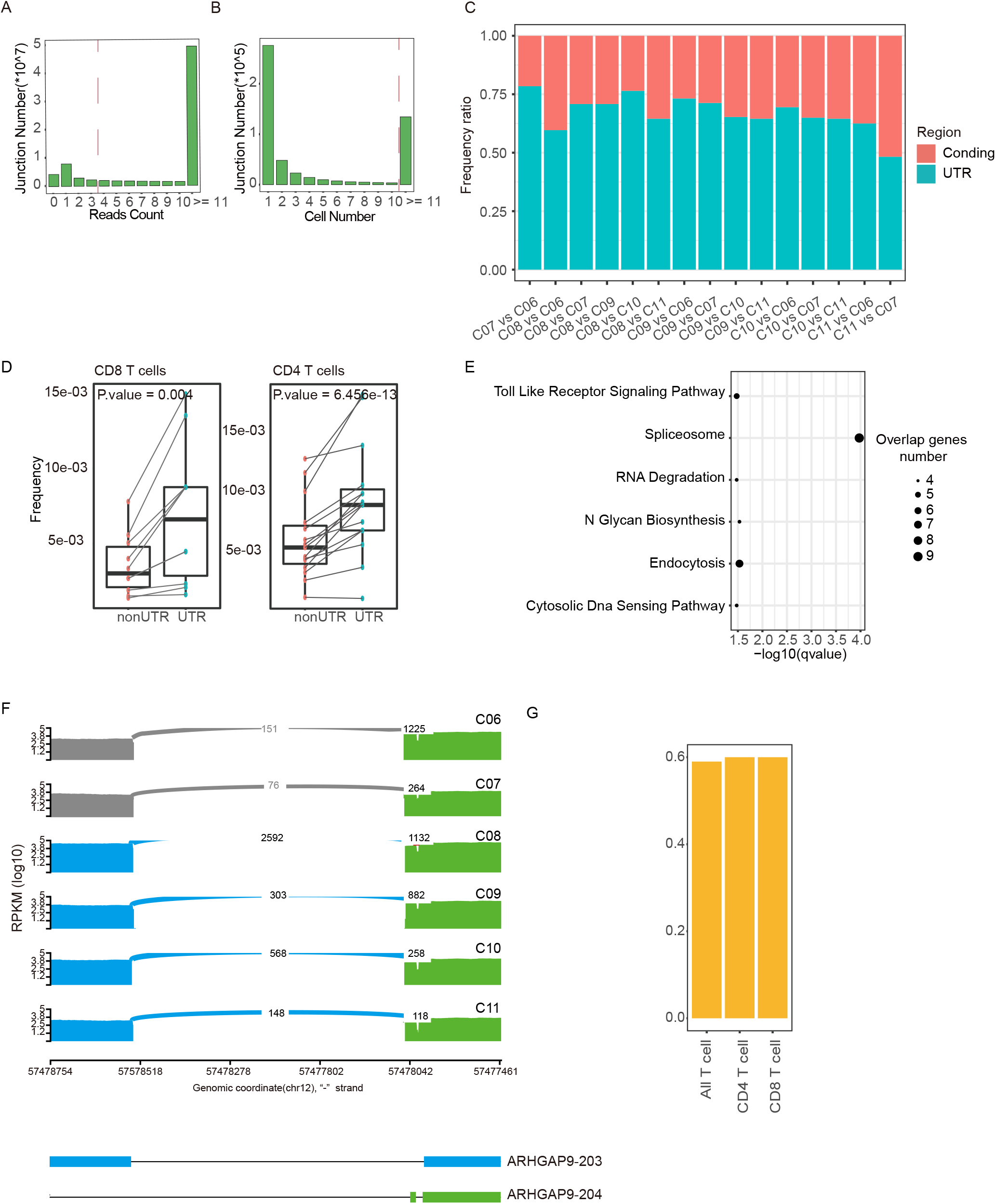
Wide differential usage of junctions in the UTR regions across T cell clusters. **A**, The read number distribution of all junction from 5,063 cells. **B**, The cell number distribution of junction with at least 4 reads supporting. **C**, The proportions of differential splicing junction in the UTR regions and the Coding regions across *CD4+* T cells. D, The frequency of DESJ in the UTR regions is significantly higher than the Coding regions. **E**, Result of KEGG pathway analysis for genes with differential splicing in the Coding regions. **F**, Sashimi plots illustrating the read distribution of *ARHGAP9in* CD4+ T cells from P0508 patient, similar to CD8+ T cells in Figure 2D. Naïve T cells (CO6_CD4.CCR7) and blood Tregs (CO7_CD4.FOXP3) show obvious differential usage of isoforms from other clusters in CD4 T cells. G, The proportion of differential expression genes with differential splicing junctions in the UTR regions.

**Figure S3.**
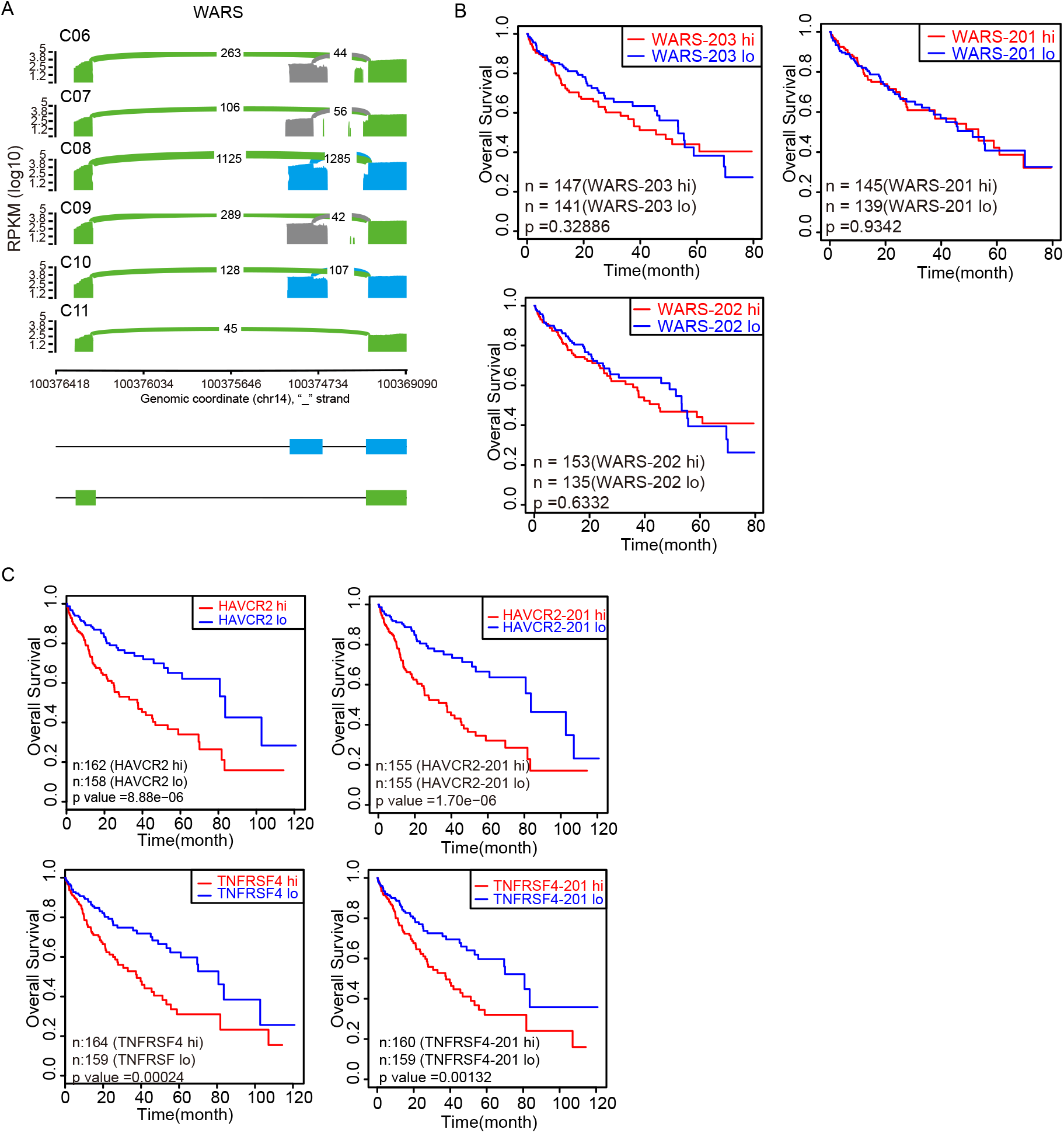
T cell heterogeneity at splicing level. **A,** Sashimi plots illustrating the read distribution of WARS in CD4+ T cells from P0508. WARS-204 highly expresses in exhausted T clusters (C10_CD4.CXCL13) and Tumor Tregs (CO8_CD4.CTLA4). **B**, Result of GO enrichment analysis for genes in Figure 3B. **C**, Disease-free-survival (DFS) curve comparing the high and low expression of WARS-201, WARS-202, and WARS-203 based on the TOGA HCC cohort. **D**, Disease-free-survival (DFS) curve comparing the high and low expression of *HAVCR2,* HAVCR2-201, *TNFRSF4* and TNFRSF4-201 based on the TCGA HCC cohort.

**Figure S4.**
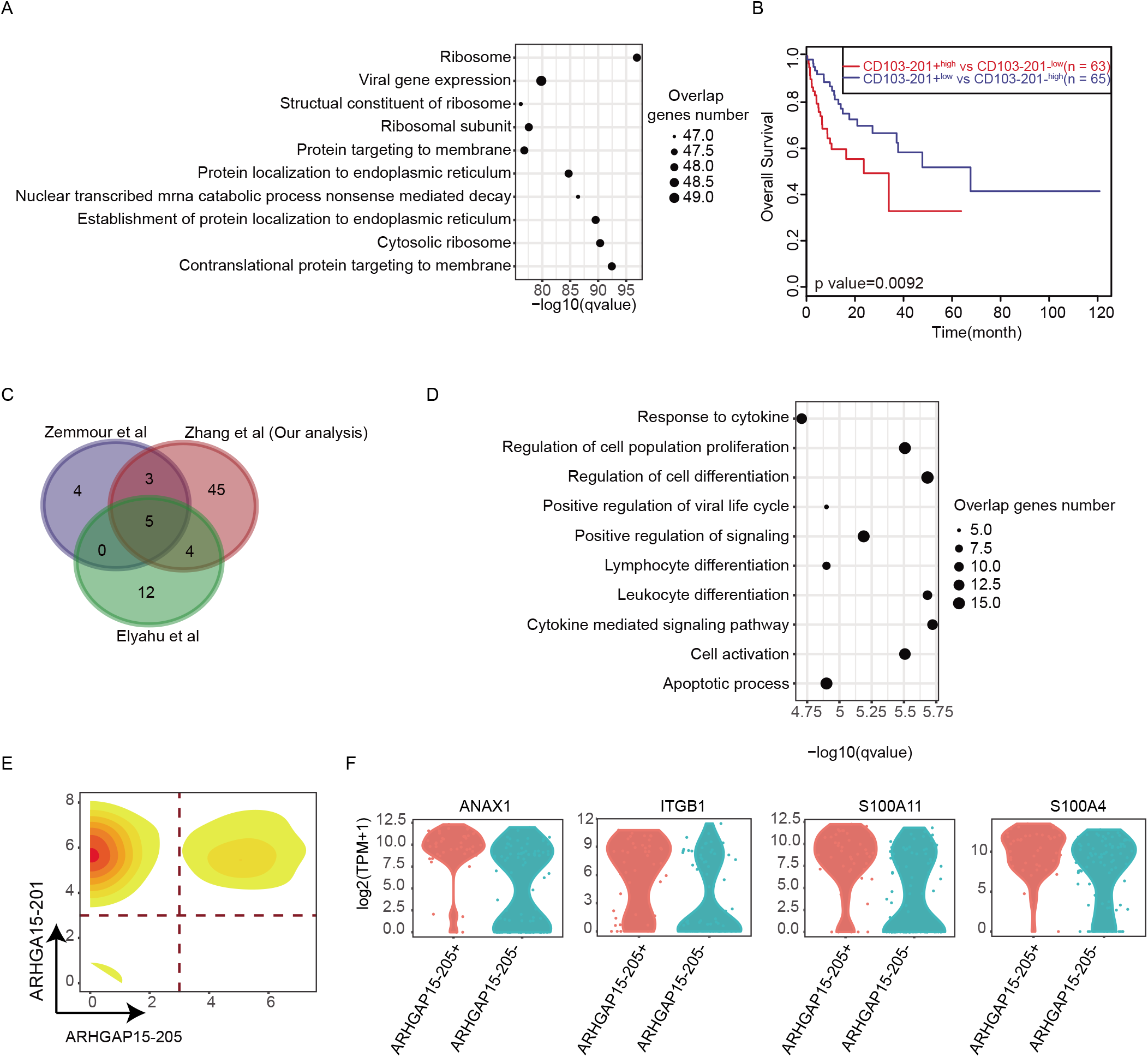
Two novel functional subpopulations identified by CD103-201 and ARHGAP15-205. **A,** Left: CD8^+^ T cells (excluding MAIT cells) was ordered along pseudotime in a two-dimensional state-space defined by Monocle2. Cell orders are inferred from the expression of DEGs across CD8^+^T cell populations. Each point with different colors corresponds to individual cells in different clusters. The middle plot shows the order of CD103-201^+^ population and CD103-201- population. Right: The exhaustion score calculated by the mean expression of gene sets related to exhaustion status (see Methods) correlated with Monocle components. Violin plots in the top corners show the distribution of exhaustion scores in various cell clusters. Different colors represent different clusters. P values were calculated by Pearson correlation, and P < 2.2 × 10-16 represents a P value approaching 0. **B**, Similar to Figure S4 A, the same pseudotime plot for four clusters of CD4+ T helper cells and the similar naiveness score calculated in all CD4^+^ T helper cells. **C**, Similar to Figure 4G, the bimodal distribution of ARHGAP15-205 shows the Intrinsic heterogeneity in naive T cells (C01_CD8.LEF1). **D**, Violin plots show the expression difference in ARHAGP15-205- and ARHGAP15-205^+^ C01_CD8.LEF1 subpopulation about activation markers, including *S100A4, S100A11, ITGB1 andANAX1.*

